# Lexical landscapes as large *in silico* data for examining advanced properties of fitness landscapes

**DOI:** 10.1101/640151

**Authors:** Victor A. Meszaros, Miles D. Miller-Dickson, C. Brandon Ogbunugafor

## Abstract

*In silico* approaches have served a central role in the development of evolutionary theory for generations. This especially applies to the concept of the fitness landscape, one of the most important abstractions in evolutionary genetics, and one which has benefited from the presence of large empirical data sets only in the last decade or so. In this study, we propose a method that allows us to generate enormous data sets that walk the line between *in silico* and empirical: word usage frequencies as catalogued by the Google ngram corpora. These data can be codified or analogized in terms of a multidimensional empirical fitness landscape towards the examination of advanced concepts—adaptive landscape by environment interactions, clonal competition, higher-order epistasis and countless others. We argue that the greater *Lexical Landscapes* approach can serve as a platform that offers an astronomical number of fitness landscapes for exploration (at least) or theoretical formalism (potentially) in evolutionary biology.

---

Historically, theoretical population genetics has often progressed by using contrived or even artificial systems and models to recapitulate the properties of natural, real-world populations of organisms[1, 2]. The fitness landscape is one of the triumphs of theoretical population genetics, an abstraction that changed how evolutionary biologists studied the process of adaptive evolution [3], and one whose original iterations were entirely theoretical. While these models are always engineered with a set of necessary constraints, they have been central to the growth of theory in modern population genetics[4].

In recent years, fitness landscapes have benefited from advances that enabled the use of empirical data towards the construction of empirical fitness landscapes[5]. Combinatorial data sets—where suites of mutations are engineered in every possible permutation—are the gold standard for these types of studies [6]. They were critical for introducing the concept of the adaptive trajectory, and have since been used as an innovation space for methods to detect higher-order epistasis [7], for metrics to calculate the speed of adaptive evolution [8], and for more rigorous attempts at predicting evolution. The limitations of combinatorial data sets are that they tend to only focus on suites of mutations within a single gene of interest, and that there are relatively few such data sets in existence [9–11]. Regardless of source, fitness measurements for these landscapes are often taken in a small number of environments, which limits our understanding of how the effect of mutations might be affected by environments. Note that this is even a problem for existing studies featuring simulated and in silico fitness landscapes [12, 13].

In this study, we propose an instrument—*Lexical Landscapes*—for generating theoretical fitness landscapes, that is not based on a algorithm. Instead, it is built on an open-source, well-vetted and established data set that can be easily analogized as fodder for the study of advanced topics in theoretical population genetics: Google Books ngram data corpora. In a prior study, this concept was introduced as a model for evolution through “protein space,” as it builds on a highly effective analogy authored by John Maynard Smith [14, 15], and can serve as a means of teaching and communicating concepts in evolutionary genetics as well. Here we double-down on this idea by arguing that the utility of the Google Books ngram corpora is not only pedagogical but also exploratory and scientific: one can use this data set to test and generate hypotheses, and develop theory in modern evolutionary genetics that exceeds the reach of current data sets (empirical and simulated). Even more, the accessibility of the data set, and connection to common words makes it easy to codify, discuss, and cross-reference.

We first outline the specific data science and computational methods necessary to generate a set of *Lexical Landscapes* of a certain kind. We then demonstrate the utility of these sets by exploring how *Lexical Landscapes* can elegantly recapitulate several standard and advanced properties of evolutionary genetics on combinatorial data sets such as fitness landscape topography and the accessibility of trajectories. For example, we introduce an “environmental context” analogy to these data, which allows us to rigorously compute properties of fitness landscapes across environments. We then move onto the elusive concept of higher-order epistasis, and examine how it is affected by environmental context. Summarizing, we re-emphasize the breadth of concepts that can be explored with *Lexical Landscapes* and speculate on its untapped potential as an instrument for modern theoretical biology.

## METHODS

### Using Google ngram values to generate empirical fitness landscapes: conceptual challenges

A prior study introduced the use of Google ngram data as a pedagogical and communicative tool for evolutionary genetics [15]. In this study, we expand this idea and argue that the Google ngram corpora has utility for thinking about more advanced concepts in evolutionary biology. To effectively utilize *Lexical Landscapes*, it is critical that one potentially confusing idea is fully clarified: while many of the patterns we observe in the data set might be reflective of culturomic [16] or evolutionary linguistic phenomenon, *Lexical Landscapes* are not engineered to study the evolution of language. Rather, they offer a transparent, open-access reservoir of data that can be easily translated into a form similar in structure to other biological and *in silico* data sets used to generate fitness landscapes. We first outline a method for collecting and curating these data (Figure 1). We then put these data to use through various calculations and simulations, which highlight the cutting-edge evolutionary questions that can be interrogated with this data set.

**Figure 1:**
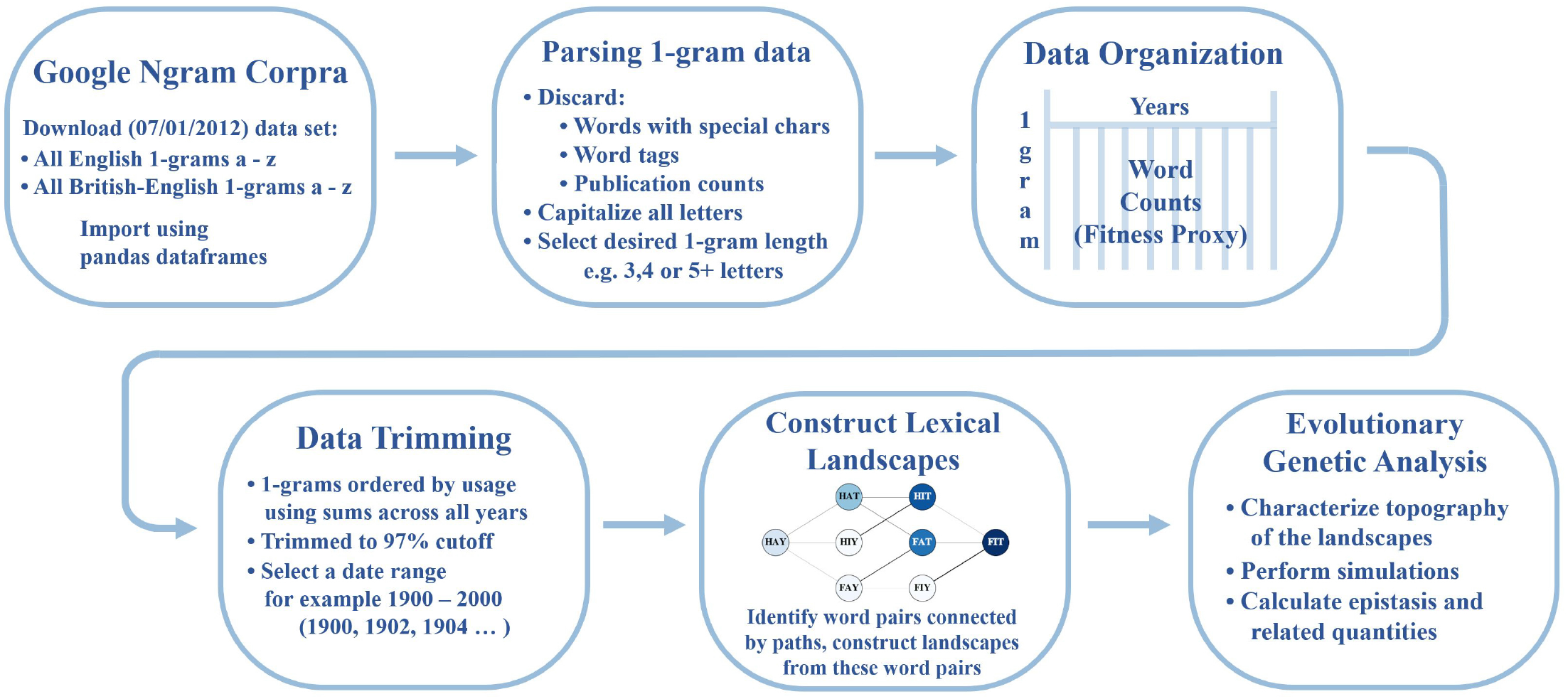
Methods Flowchart. The flowchart above describes the methodology used to construct *Lexical Landscapes* and carry out several of the additional analyses discussed in this study.

### Data acquisition and curation

The strength of the *Lexical Landscape* approach resides in its expansiveness. Given the *English* alphabet of 26 letters, there are over 600 different 2-letter “alleles” (different word variants) and over 17,500 3-letter alleles. While the majority of words will have near-zero usage frequencies, or what we call *Lexical Fitness*, there might be utility in studying a large portion of these landscapes. Thus, we propose that the innovation of this approach resides in its ability to create an enormous number of non-arbitrary fitness landscapes. In order to demonstrate how this data can be leveraged for evolutionary theory, we have focused on combinatorial and empirical landscapes of smaller size. A fuller description of the size of the *Lexical Landscape* data set (actual and curated) can be found in the Supplementary Appendix.

To construct the data set of *Lexical Fitnesses*, we formatted data provided in Google’s ngram corpora. Specifically, we downloaded the entire *English* corpora of *1-grams*—or single words, as opposed to 2-grams for instance, which are pairs of words—for all the letters in the alphabet. We discarded 1-gram data sets containing numbers or special characters. The data from Google’s corpora originally included “word tags” (e.g.__NOUN__, __VERB__, __ADJ__) specifying the grammatical context in which the word was used—this provided a breakdown of how often a given word was used as a noun, adjective, etc. In our data set, we simply removed these tags and ignored their grammatical context. The word counts associated with each word are therefore the *total* usage counts, which are sums over these grammatical contexts. Lastly, the original data includes counts enumerating the number of books each word appeared in; these were also discarded for our purposes (although, this information and others that are available could be useful for other purposes).

For this study, the data is composed of all 3, 4, and 5-letter words in Google’s *English* corpora, along with the number of times each word was used in every other year beginning in 1900 and ending in 2000. The ngram word frequencies are taken from books that Google has been able to survey (approximately 6% of all books ever published [17]). We limited the data to every other year to expedite computations and we chose to use the 20th century data as it represents the most modern full century from the Google corpora—it is also the most densely populated in terms of word usage. Note, however, that the Google ngram corpora data go back to the 1500s, and so there is nothing preventing the construction of *Lexical Landscapes* for any of these years.

**Table 1:** Defining concepts and terminology in Lexical Landscapes

| Concept | Definition | Example |
| --- | --- | --- |
| • Fitness landscape<br>• Adaptive landscape<br>• Fitness Graph | These terms are used to describe a kind of genotype-phenotype map that organizes information in a manner that communicates details about the evolutionary process. | See Figure 2 |
| • Accessible pathway or trajectory | An evolutionary pathway through a fitness landscape where consecutive step leads to an increase in the fitness proxy. | See Figure 3 |
| • Within path competition ($C_w$) | Across a given accessible pathway, there is competition between alleles. Recent work has demonstrated that the this total competition is powerfully associated with the speed of evolution across a given pathway | See Figure 5 |
| • Environment or context | How evolution occurs is profoundly influenced by its environmental context. In *Lexical Landscapes*, either the time dimension (continuous variable) or language variable (categorical) can be used as an analogy for environment or context. | Time or language.<br>For example, the CARS → SOME landscape can be constructed at any of the available time points in which there are data. In Figure 2, we construct landscapes |
| • Genotypic Context | How a fitness landscape for a given gene is affected by the whole-organism genome in which it is embedded. For example, in transgenic studies where a gene from organism A is engineered into the genome of organism B. It is possible that this change in genomic context can change the phenotypic effects that gene. In this study, we use British-English as an analogy. The CARS → SOME landscape features can be calculated in certain sub-“genera” of English, like British-English. | See Supplementary Appendix |

We divided the data into the three word-lengths—3, 4, and 5—and within each we ordered the words according to word popularity. More precisely, for each word within each word-length category, we summed the total word count usage for that word across all the years in our data set—a total of 51 years: 1900, 1902,…, 1998, 2000. Words with a higher sum were assigned a higher popularity. For instance, we found that the most popular 3, 4 and 5-letter words—calculated in this manner—were ‘THE’, ‘THAT’, and ‘WHICH’, respectively. In order to use words that are fairly popular and to avoid acronyms and initialisms—also to facilitate quicker calculations—we truncated each list of words down to the top 200 most popular 3-letter words, the top 1500 most popular 4-letter words, and the top 5,000 most popular 5-letter words. The popularity cutoffs were chosen so that the sum of all word counts in each truncated list represented roughly 97% of the total summed usage-count of all words in that word-length category. We chose 97% so that the data set was as large as possible without including many of the very numerous, though infrequently used, acronyms/initialisms.

Within each truncated list, we identified various *word-paths*. A word-path is characterized by a starting and an ending word and comprises a collection of words that take one from the beginning word to the ending word by way of changing one letter at a time. More precisely, a letter-change occurring at some location in the word is a swap of a letter in the beginning word for the corresponding letter in the ending word, such that at each step the combination of letters is a word itself. An example can be seen in Figure 3. We will call the collection of *all* letter combinations between two terminal words a *landscape*—we will analogize this to a *fitness* landscape—and two examples of which are shown in Figure 2. Word-paths were interesting to consider since the words that form a path each have some kind of lexical significance, and can be analogized with *genotypes* with sufficiently high fitness. Whereas, arbitrary combinations of letters—which do not make good *lexical* sense—are analogized with genotypes that do not fair well in biological settings, having low fitness (i.e. they do not make good *biological* sense). Using word-usage frequency as a proxy for fitness, we identify among the set of word-paths those which are “evolutionarily favorable”—i.e. those for which the fitness value increases along the path, in a given year. We refer to these paths as *accessible* or sometimes *uphill* paths. The details of the algorithm used to identify these paths, and how to access the Python script, are described in the Supplementary Appendix. It is very important to note that even with the selective criteria used to curate the greater ngram corpora, we can still generate over 1 million total fitness landscapes, that is, combinatorial sets with Lexical Fitness values between pairs of alleles. This would constitute, by many measures, the largest set of fitness landscapes in existence.

**Figure 2:**
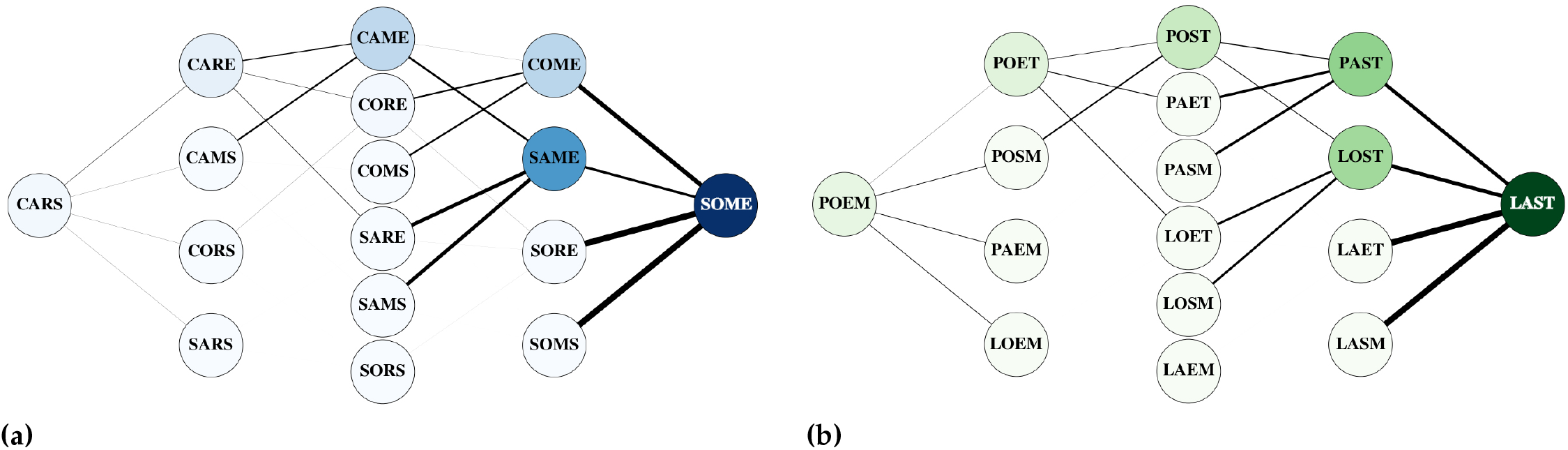
Fitness graphs for the CARS → SOME and POEM → LAST landscapes. Nodes are connected if they differ by a single letter. Darker nodes represent higher fitness values, and edges are weighted by the difference in the fitness values (in absolute value).

**Figure 3:**
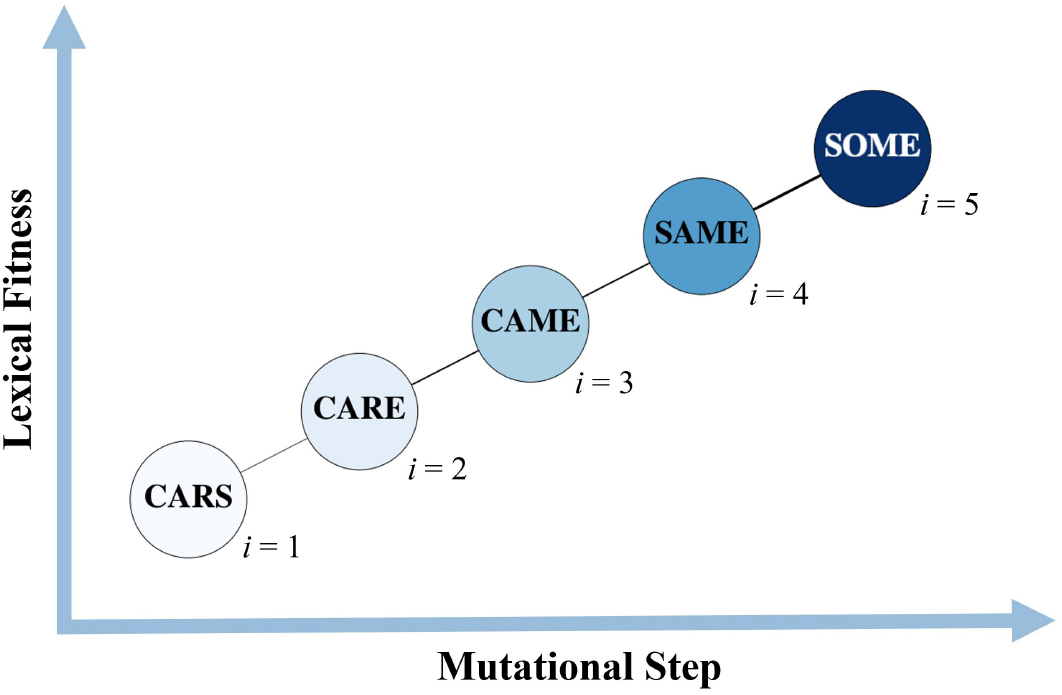
Adaptive trajectory. A hypothetical “uphill” word-trajectory in the CARS → SOME landscape. Each word is indexed by *i*, referenced in eq. 1, and the color gradient indicates the fitness value *r_i_*, with darker blues showing higher fitness. The edges connecting the nodes are thicker and thinner, depending on the difference in the fitness values (in absolute value).

### Choice of example Lexical Landscapes used for illustrative purposes

In order to examine various properties of fitness landscapes using *Lexical Landscapes*, we chose two model landscapes in the set of 4-letter words, CARS → SOME and POEM → LAST. These two were chosen among thousands of possible landscapes because they carried features that make them useful examples:

- Their topography changed with environmental contexts, which made it likely that evolutionary dynamics might differ across context.
- Their topography is rugged, indicating the presence of epistasis. Rugged landscapes often create non-intuitive evolutionary patterns.
- They contained multiple paths that were accessible across several environments. These are also features of landscapes that are likely to offer non-intuitive dynamics, as these landscapes possess multiple accessible paths which can *compete* in an evolutionary sense.

Note that none of these above outlined features of model landscapes are essential for the use of *Lexical Landscapes* as an instrument for studying and exploring properties of fitness landscapes. We only imposed these criteria on the landscapes in order to illustrate the potential utility of this tool. Any one of thousands of Lexical Landscapes might be generated for the study of advanced properties of fitness landscapes. In the Supplementary Appendix, we provide similar analyses for 3 and 5-letter word *Lexical Landscapes*.

In what follows, we show how three concepts in evolutionary biology can be explored with this approach. We first consider how fitness values can be used as a proxy for growth rates in order to simulate the “growth” and evolution of the various alleles, or word combinations, that comprise a landscape. We compare simulated dynamics in various contexts. Second, we explore how lexical fitness values can be used to construct a metric for the expected *time* associated with evolution along a particular trajectory or *word-path*—we refer to this metric as the within-path competition. Lastly, we use lexical fitness to calculate the degree of epistasis present in the data set. We observe how this degree may vary over years across the 20th century.

### Evolutionary dynamics with the Discrete Asexual Reproductive Population Simulator (DARPS)

Here we discuss the simulated evolutionary dynamics for the two example 4-letter landscapes (CARS → SOME and POEM → LAST). The purpose is to illustrate how simulations across *Lexical Landscapes* can demonstrate many properties, simple and advanced, that are direct analogues to biological processes.

Each landscape consists of the sixteen letter combinations for a given word pair, featuring all possible letter-swaps between the words in the word pair (Figure 2). In the simulation, the *population* of the first word (allele) in the word-pair was set to 1000, while all other words had an initial population of 0. Each population was allowed to evolve at each time step according to some fixed probability of mutation (we chose 10^−8^), a mutation rate that is on the order of what we observe in microbial populations. Each word was also assigned a growth rate which was correlated with its *Lexical Fitness* for a given year in the following way: we first calculated the average fitness value of all words in the landscape for the given year. Then, we use the fitness values in the landscape divided by this average for the given year as the growth rates for each word. Note that the growth rates were fixed throughout the simulation and only depended on the fitness values for the year chosen. In this way, the growth rates are defined by their *relative* fitness (relative to the mean fitness of all words in a given landscape for a given year). We used the *Discrete Asexually Reproducing Population Simulator* (DARPS), a simulator of evolution in large populations of organisms that resembles microbial populations [8, 18]. At each time step in the simulation, or generation, a certain proportion of each word’s population undergoes replication (in the generic exponential sense, with the growth rate as the Malthusian parameter). Mutations occur during replication according to the probability of mutation. As the simulation progresses, different alleles can rise and fall in frequency. We can visualize the dynamics of these simulations by graphing the fraction of each allele in the population (Figure 4).

**Figure 4:**
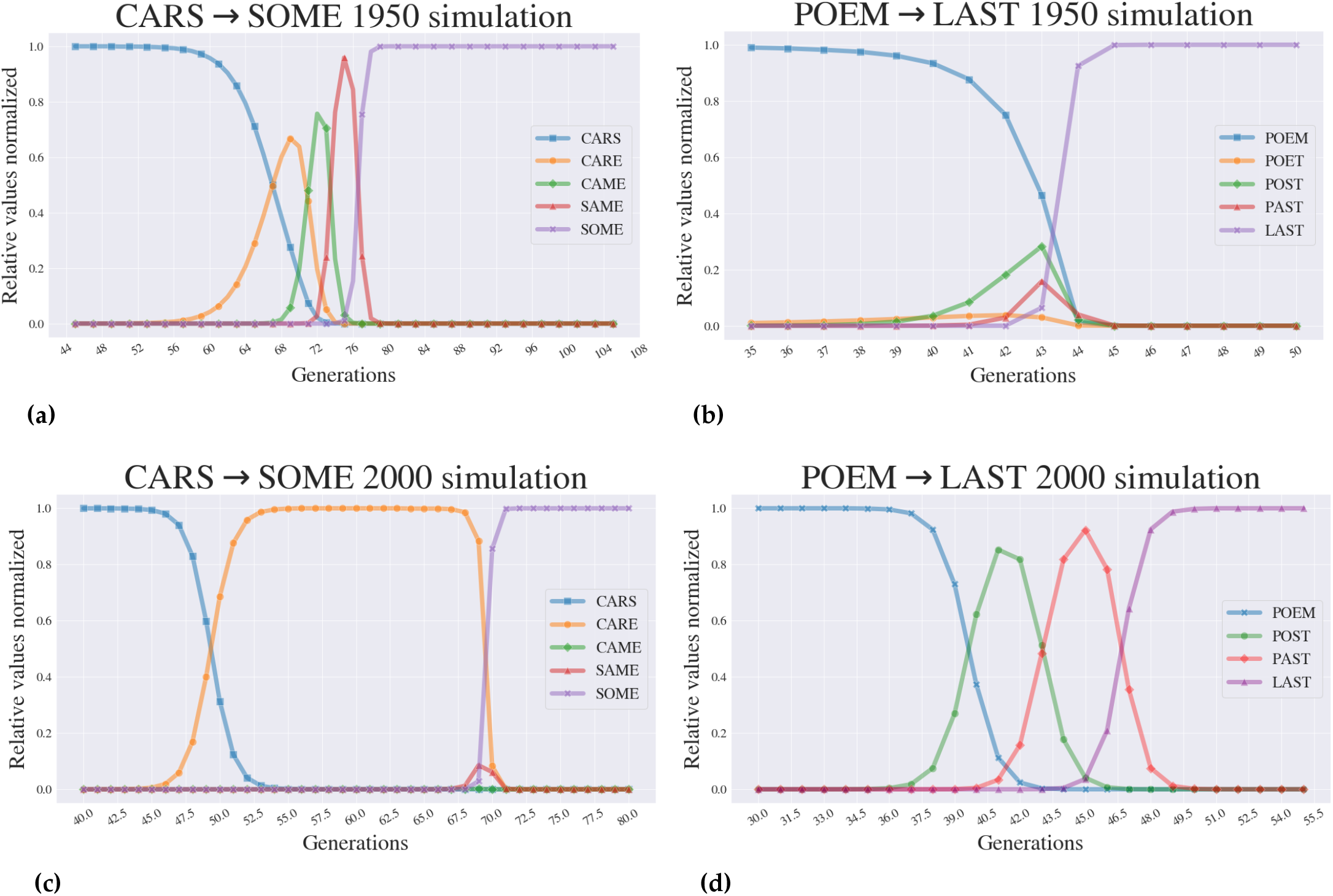
Simulated dynamics across different contexts demonstrate different dynamic properties. Figures (a) and (c) show illustrative simulations across the CARS → SOME *Lexical Landscape*. Figures (b) and (d) show two for the POEM → LAST simulation dynamics. The y-axis shows the percentage of the population occupied by a given word. All simulations begin with 100% of the population fixed at the 0000 genotype and ultimately reach fixation at the 1111 genotype. Note, however, that the dynamics through which this occurs changes as a function of context, or year in our case.

### *Within-path competition* and the speed of adaptive evolution across a fitness landscape

We now consider how *Lexical Landscapes* can be used to examine advanced concepts that have never before been explored at the scale that *Lexical Landscapes* offers. Recent studies [8] have introduced a term that correlates powerfully with the time associated with evolution across a landscape and is calculated for a given *path*. It is termed the *within-path* competition (*C_W_*) and it is defined by the formula,

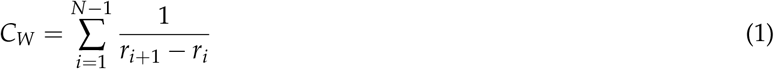

where *r_i_* is the *growth rate* of the *ith* allele, and where *N* is the length of the evolutionary-path (the number alleles in the path)—*C_W_* can therefore be thought of as a sum over the *links* (or edges in a graph, such as in Fig. 3) in a path. *C_W_* is typically computed using growth rate measurements, however, in this work we use the *Lexical Fitness* values as a proxy for growth rate. When *C_W_* is high, evolution across a trajectory is expected to be slow, and evolution is fast when *C_W_* is low, provided that *r*_*i*+1_ > *r_i_* for each *i*, which assumes that the path is an *uphill* path. In the context of *Lexical Landscapes*, we calculate *C_W_* along *word-paths* in the landscape. These paths are analogous to a series of mutational steps in a 4 loci allele, progressing along an evolutionary path, such as 0000 → 0001 → 0011 → 0111 → 1111. Figure 5 shows two such word-paths, one in each of our example landscapes. One can observe how drastically this value can vary across time. We present more examples of *C_W_* in the Supplemental Appendix.

**Figure 5:**
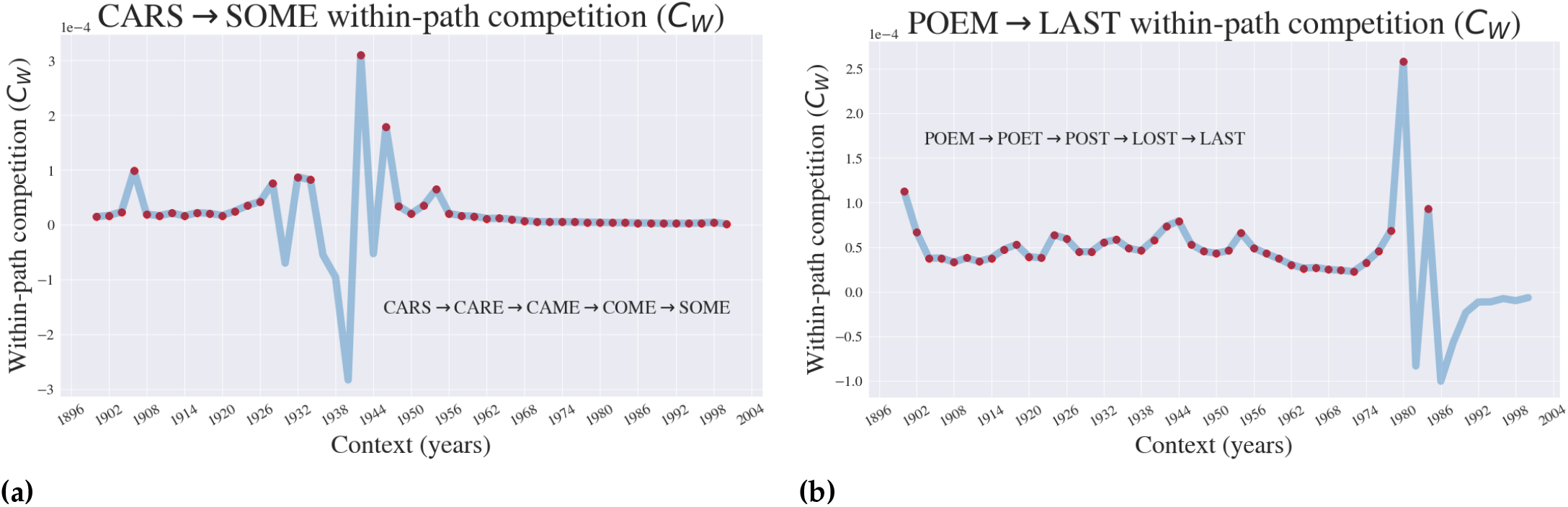
Within-path competition (*C_W_*), a proxy for the speed of evolution changes with context. The *within-path* competition coefficient is shown for specific paths (shown in the figures) in the CARS → SOME (a) and POEM → LAST (b) landscapes. Red dots indicate the years for which *C_W_* was calculated on an *uphill* path, which is to say that the associated path had increasing fitness values as the path is traversed.

### Calculating higher-order epistasis

*Lexical Landscapes* also offer a way to consider theoretical questions related to the subject of epistasis within a landscape. In particular, one may consider how the epistasis manifests across different environmental contexts.

Epistasis remains a cutting-edge topic in evolutionary biology that continues to be the object of study for a variety of reasons, and measured using diverse methods[10, 19–21]. For our purposes, we use a Walsh-Hadamard transformation of the fitness values, scaled by an additional diagonal matrix, as presented in Poelwijk et al. [19]. We summarize the approach below.

For a given year, the *Lexical Fitness* values for each letter combination in a given landscape (CARS → SOME or POEM → LAST) are arranged into a vector *x*—for 4-letter words there are 16 fitness values, one for each letter combination in the landscape. In short, this vector will be multiplied by a 16 x 16 square matrix; we then take the absolute value of the output and normalize. The 16 x 16 matrix is the product of a diagonal matrix *V* and a Hadamard matrix *H*.

These matrices are defined recursively by,

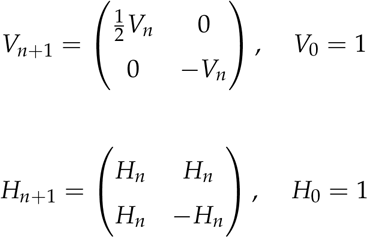

where *n* is the number of loci (*n* = 4 for our purposes as we consider 4-letter words). The output *y* of this matrix multiplication is given explicitly by,

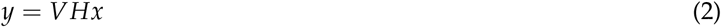

where *V* and *H* are the matrices above for *n* = 4. We then take the absolute value of the entries in *y* and divide each entry by the sum of the absolute values to normalize. In this paper, we refer to these as the *epistatic* coefficients *E_i_*.

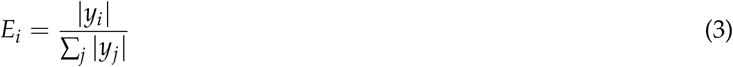

The absolute value and normalization was performed in order to focus exclusively on the magnitude of the epistatic effects. We will sometimes use bit strings such as ‘0101’ as the index *i* when referring to the epistatic coefficients *E_i_*. For instance, *E*_0101_ represents a measure of the epistatic effect of “mutations” in the 2nd and 4th letter, or locus.

In addition, one may consider averaging these epistatic coefficients *E_i_* within what we call an *order*, enabling comparisons between orders. An *order* is the collection of mutations of an allele with the same *number* of mutations, or bit-flips (or letter *swaps* in a given combination of letters in the landscape). For instance, the coefficient *E*_0000_ belongs to what we refer to as the “0th” order (zero letter swaps), whereas *E*_0001_, *E*_0010_, *E*_0100_, and *E*_1000_ all belong to the “1st” order (one letter swap), etc. Note that, like the 0th order, there is only one coefficient contained in the 4th order, namely *E*_1111_. Taking the average of the epistatic coefficients within each order presents information about the general epistatic effect based solely on the number (or order) of mutations (letter swaps) in a given landscape, one swap, two swaps etc. In Figure 6, we present these average epistatic effects for our two landscapes (we label them with the term “absolute mean” since we incorporated absolute values and averages in the calculation). In the Supplemental Appendix, we show all the epistatic orders (without averaging) for our two case landscapes, as well as two additional ones within the 3 and 5-letter categories—in the Supplemental Appendix, we refer to these plots as the *dis-aggregated* epistasis, in contrast to the *aggregated* epistasis we present here in the main text. As past studies have focused on comparing higher-order effects [7], one can glean a lot of information from aggregating all effects by their order. For example, we can observe whether a given landscape is dominated by epistatic effects of a certain order, and speculate as to why this is so.

**Figure 6:**
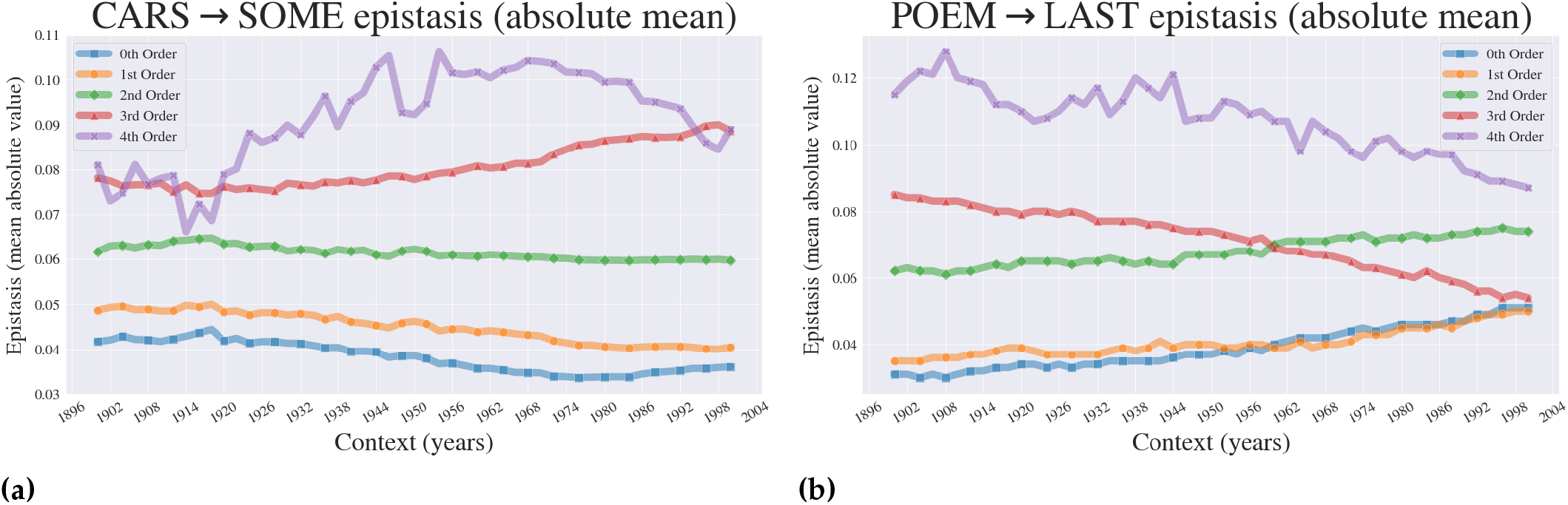
Higher-order epistasis across context. Figures (a) and (b) show how epistatic effects for our example word landscapes can vary across environment, or in this case across time.

## RESULTS

### Simulations of evolution across *Lexical Landscapes*: standard and non-standard dynamics

Figure 4 demonstrates that *Lexical Landscapes* can be used for the construction of simulated adaptive evolution as observed in several studies of evolution across empirical fitness landscapes [5, 6, 22, 23]. The *Lexical Landscapes* used as models in this study also demonstrate advanced properties of evolution. While Figure 4a displays the standard step-wise evolutionary trajectory whereby the population reaches appreciable frequencies at all alleles in a given path, Figures 4b-d demonstrate how certain simulations display features of *stochastic tunneling*, where evolution appears to “skip steps”. That is, when an intermediate genotype makes no appreciable appearance in population space. Prior studies have revealed that stochastic tunneling can happen when populations sizes are large and mutations rates are high. [24, 25].

### The accessibility of pathways and within-path competition (Cw; a proxy for the speed of evolution) changes as a function of context

Figure 5 shows how one may consider visualizing the *time* that evolution takes within a landscape across varying contexts. This translates to the idea that the speed of adaptive evolution is a function of the environment that it’s in. In the Supplemental Appendix, we compare the plots in figure 5 to two other paths within each landscape, in addition to considering *C_W_* for other word lengths.

### The magnitude of higher-order epistasis changes across environmental contexts (years)

Next, we turned our attention to a popular concept in evolutionary genetics: higher-order epistasis. Using the Walsh-Hadamard transformation (see: Methods), we calculated the aggregated higher-order epistasis for each order (0th - 4th) and plotted it across the contexts from 1900–2000. Strikingly, we observe powerful context dependence of higher-order epistatic effects (Figure 6), for both examined landscapes (CARS → SOME and POEM → LAST). Interestingly, we can also see that across most of the queried environmental contexts (1900 - 2000), fourth order effects are predominant, indicating that within these landscapes, words derive their utility (in terms of word-frequency, or *Lexical Fitness*) from interactions between all four of the letters. One also notices that the predominant epistatic influence across the landscape changes across contexts. There are, for example, small contextual windows (years) in the CARS → SOME landscape where the 3rd order effects dominate. Keep in mind that the analysis in Figure 6 obfuscates the direction of epistasis: as outlined in the methods, absolute values of the output vectors were computed, which means that information on the sign of individual effects is lost. The data for the entire calculation, however, can be found on GitHub: github.com/ogplexus/LexicalLandscapes

## DISCUSSION

In this study, we introduce a method for creating a database for a large number of empirical fitness landscapes based on the ngram corpora. Though these data are not “biological,” they are empirical in the sense that the data for the individual nodes arise from a measurement of a natural system such as a language, and are not simulated. For this reason, *Lexical Landscapes* provide an ideal hybrid empirical-*in silico* data set for readily generated fitness landscapes at a size and scope that far exceed that of current technology-limited biological systems.

We also demonstrate how to put these data to use in a way that highlights how one can observe advanced and higher-order phenomena in evolutionary genetics. Importantly, we emphasize that a strength of the *Lexical Landscape* is in its capacity to provide analogies for varying environmental contexts, using time as one such means for variation. We do so using three different exercises.

Firstly, we demonstrate how one can observe evolutionary simulations across these landscapes. These simulations follow the step-wise process of adaptive evolution that has been observed for similarly constructed empirical fitness landscapes. We also observe, however, circumstances where evolution displays non-standard properties. Specifically, the topography of some landscapes displays features of “stochastic tunneling,” where a population appears to “skip steps” in adaptive evolution. The reality is that evolution isn’t “skipping” steps at all, but rather, that an intermediate allele is present in low frequency, but due to high overall population size and mutation rate, subsequent steps are traversed.

Next, we examined how within-path competition (Cw), a measure of the clonal competition acting along a particular adaptive trajectory. This has been demonstrated to be strongly correlated with the speed of evolution across a certain trajectory[8], and so this analysis offers a demonstration of how one can gain a picture for the predicted speed of adaptation as a function of a fitness landscape occupying some specific niche.

Lastly, and most provocatively, we demonstrate how the magnitude of higher-order epistasis is altered by context. Despite the fact that epistasis is a powerfully controversial topic, very few studies have explored how context influences higher-order epistasis [26–28]. Using *Lexical Landscapes*, we are easily able to generate fitness landscapes, calculate epistatic coefficients within single environments, and demonstrate how those coefficients manifest across a wide number of environments. This is a understudied phenomenon, one for which no general theory or rules have yet been constructed. Using *Lexical Landscapes*, we now have insight that can be applied to existing biological data. There are many reasons why more careful interrogation of how epistatic effects change with environment might be relevant, as it could explain difficulties in recovering individual SNPs of large effect in large-scale genomic studies.

The empirical*-in silico* dichotomy is one that has been a part of population and evolutionary genetics since its inception, and will always have a place. Certain questions will always benefit from volumes of data that are beyond the scope of what the biological world offers. With *Lexical Landscapes*, we offer a method for generating data sets that can be used to explore many features of adaptive evolution.

## SUPPLEMENTARY INFORMATION

The Supplemental Appendix contains a number of important explanations and extensions of the *Lexical Landscape* approach. They include:

- A description of the size of the data set utilized in this manuscript
- An examination of how Lexical Landscapes can be explored in other language subsets (e.g. British English)
- Further details and calculations of higher-order epstasis
- Added investigations into landscapes of 3 and 5 letter 1grams

## DATA AVAILABILITY

Code and data sets featured in this manuscript can be found on Github: https://github.com/OgPlexus/Lexical.Landscapes

## ACKNOWLEDGMENTS

The authors would like to thank D. Weinreich, A. Alexander, and S.J. Gates for helpful discussion and input on the project. CBO acknowledges funding support from NSF RII Track-2 FEC (Award Number: 1736253), “Using Biophysical Protein Models to Map Genetic Variation to Phenotypes.”

