## Supplemental Appendix for "Lexical landscapes as large *in silico* data for examining advanced properties of fitness landscapes"

In this section, we provide additional information and further analysis that highlight features of the *Lexical Landscapes* approach that exceed that which were described in the main text. The main text was dedicated to outlining the origins, methodology and several simple applications of *Lexical Landscapes*. It should not be interpreted as representing the full capacity of the method, as thousands of *Lexical Landscapes* can be created of various size and topography. In addition, we provide more detailed analysis of higher-order epistasis for the examples discussed in the main text. Lastly, we demonstrate how similar these calculations are to real-world calculations on empirical fitness landscapes.

#### *Lexical Landscapes* databases

| | Available words | Possible landscapes |
|---|---|---|
| **3-letter** | 17,338 | 7,665,017,103 |
| **4-letter** | 163,809 | 684,247,229,136 |
| **5-letter** | 419,185 | 4,480,748,948,520 |
| **Total** | 600,332 | 5,172,661,194,759 |

**(a) All English data set**

| | Available words | Possible landscapes |
|---|---|---|
| **3-letter** | 16,375 | 6,837,168,375 |
| **4-letter** | 111,675 | 318,015,445,725 |
| **5-letter** | 267,943 | 1,830,726,174,303 |
| **Total** | 395,993 | 2,155,578,788,403 |

**(b) All British-English data set**

| | Available words | Possible landscapes |
|---|---|---|
| **3-letter** | 200 | 1,014,900 |
| **4-letter** | 1,500 | 57,336,750 |
| **5-letter** | 5,000 | 637,372,500 |
| **Total** | 6,700 | 695,724,150 |

**(c) Reduced English data set**

| | Available words | Possible landscapes |
|---|---|---|
| **3-letter** | 250 | 1,587,375 |
| **4-letter** | 500 | 6,362,250 |
| **5-letter** | 5,000 | 637,372,500 |
| **Total** | 5,750 | 645,322,125 |

**(d) Reduced British-English data set**

**Figure S1: *Lexical Landscapes* subset database sizes** Tables **(a)** and **(b)** show the number of 1-grams of each type gathered from the *All English* and *British-English* Googled Ngram Databases for this study, as well as the possible associated *Lexical Landscapes* that could be constructed from each set. Tables **(c)** and **(d)** likewise show the number of available words and the total possible landscapes that could be constructed from the reduced data sets used within this investigation. The reduced data sets were constructed using cutoffs in order to eliminate words with excessively low usage counts and avoid the presence of acronyms within *Lexical Landscapes*

In Figure S1 we present the total number of words available within the data that was gathered from the Google ngram corpora as well as the possible number of *Lexical Landscapes* which can be generated using these words. The “total data set” is divided into six subsets, three *All English* 3-letter, 4-letter, and 5-letter parcels and three *All British-English* 3-letter, 4-letter, and 5-letter parcels.

For our initial investigation, we sought to generate landscapes with features that facilitated use in studying properties of empirical fitness landscapes. This necessitated the implementation of filters against landscapes that contained excessive acronyms and words or very small usage counts. For that reason, a reduced subset of the entire full data set was used to construct our featured landscapes. The number of words and possible landscapes in these reduced sets are displayed in Figure S1. The constraints and cutoffs that we used to generate the data set are arbitrary and were only put in place in order to limit our initial search. Note that, even with the filters, we have generated well over 1 million fitness landscapes that can be studied. All the data used in this study can be found on GitHub: https://github.com/OgPlexus/Lexical.Landscapes.

Unique Lexical Landscapes are defined between pairs of words, independent of word order (e.g CARS → SOME is the same as SOME → CARS). Given some number of available words from one of the six parcels, the number of possible landscapes is determined combinatorially (n choose 2) as the number of unique pairs of words, across the 51 contexts (years from 1900 - 2000), as shown in the equation below:

$$AvailableLandscapes = [[WordCount \times (WordCount - 1)] \times Years]/2 \quad (1)$$

#### Dis-aggregated epistasis from Figure 6 in the main text

The calculated aggregated epistasis from Figure 6 in the main text represent averages over coefficients within each order; e.g. what is labeled as “1st order” is the average of the absolute values of the coefficients corresponding to 0001, 0010, 0100, and 1000. This is consistent with prior representations of higher-order epistasis using the Walsh-Hadamard Transform [1]. Alternatively the epistatic effects between *all* mutations can be observed individually as well. In the plots shown below, we show what we refer to as the *dis-aggregated* epistasis, and how it varies across context. The values plotted are the absolute values of outputs of the Walsh-Hadamard transform, discussed in the main text, and there is no averaging. Thus, instead of the 5 lines shown in the main text, one for each order: 0th, 1st, 2nd, 3rd, and 4th, there is a full set of 16, one for each output of the WH transform.

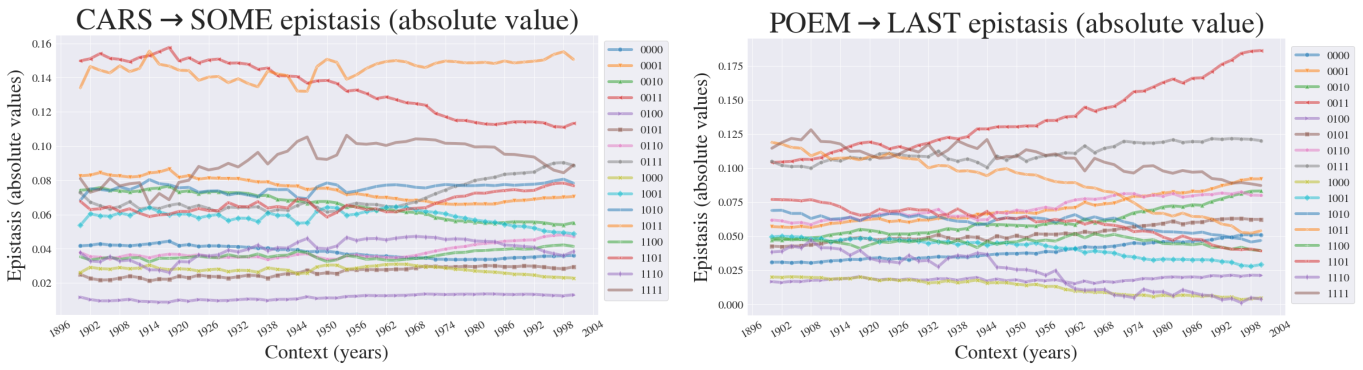


**(a)** **(b)**

**Figure S2: Dis-aggregated epistasis across Lexical Landscapes.** The main text Figure 6 demonstrated the aggregated effects, the coefficients corresponding to 0th - 4th order effects. Figures **(a)** and **(b)** show how individual epistatic effects can vary over time, or more generally, across context. Each line represents a particular epistatic effect, and the lines are grouped by order and demarcated by color theme and marker. For instance, the collection of light red to dark red lines shows the *first* order epistatic effects: 0001, 0010, 0100, 1000.

In addition to taking the absolute values of the outputs of the Walsh-Hadamard transform, we also normalize to unity at every time step/change in context. That is, the total sum of the values shown by the lines is one for each year. We choose to take the absolute value to bring attention to *magnitude* only, and normalization at each time step is performed to emphasize the *relative*/proportional differences between epistatic coefficients. Each plotted line is labeled by a bit string, representing one of the epistatic coefficients.

For instance, the line labeled “0101” shows the coefficient measuring the epistatic effect of mutations in the 2nd and 4th loci. In the figures below, we examine this dis-aggregated epistatic behavior in our choice pair of 4 letter 1-grams, featured in the main text.

#### An analysis of British-English: An analogue for exploring properties of fitness landscapes across a specific genomic background

As outlined in the main text, our primary investigation examines the characteristics of a pair of fitness landscapes constructed using four letter 1-grams as gene analogues. These four letter 1-grams were drawn from the entire Google ngram English language corpora. This complete English data set includes words from all the books published in all the diverse dialects and variants of the English language, for example British English, Scottish English, Austrian English etc. To each a variation on the other, they are all subsets of the superset that is the English language.

Similarly, an analogue can be drawn here to subset of species within the same taxonomic order. For example, the main text might have demonstrated the effects of a given suite of mutations for a protein across a particular genera. If one were interested in a specific species, however, they would have to measure that fitness landscape only in that species. These sub-sets of the English language corpora could be analogized as such subspecies. British English would be a "species" in the English "genus." We provide calculations of properties in British English to highlight how not only environmental context (analogized as time in our study), but genomic context can influence properties of fitness landscapes and evolution.

##### Aggregated epistasis - British-English

Epistasis values aggregated by order are also provided for the British-English analogues of the CARS → SOME and POEM → LAST landscapes.

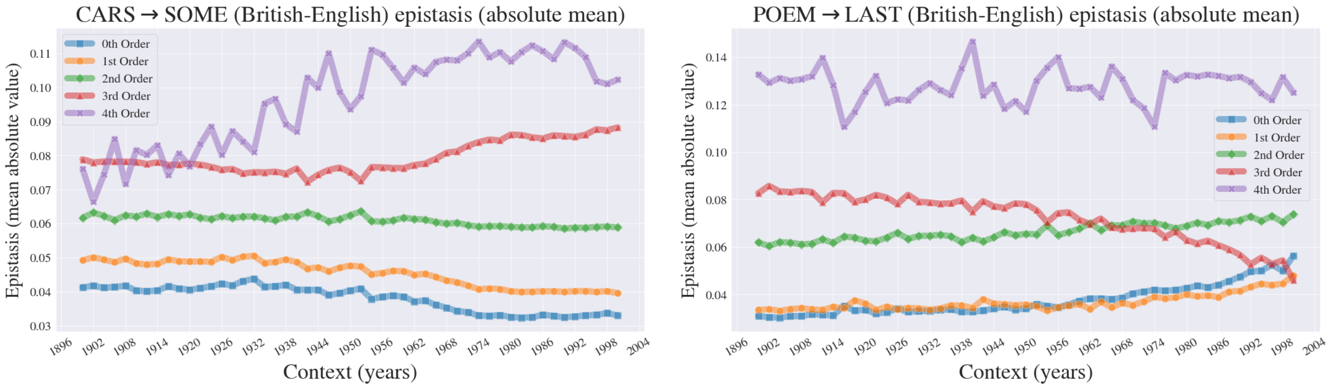


**(a)** **(b)**

**Figure S3: Aggregated British-English higher-order epistasic effects.** Fitness graphs for the POEM → LAST landscape. As described in this manuscript, the aggregated epistatic effects combine the averages of individual interactions and organize them by their order. Figure S3 shows these effects for the British-English CARS → SOME and POEM → LAST landscapes.

### Dis-aggregated epistasis - British-English

Here we show all the individual, dis-aggregated epistatic interactions for the British-English analogue landscapes.

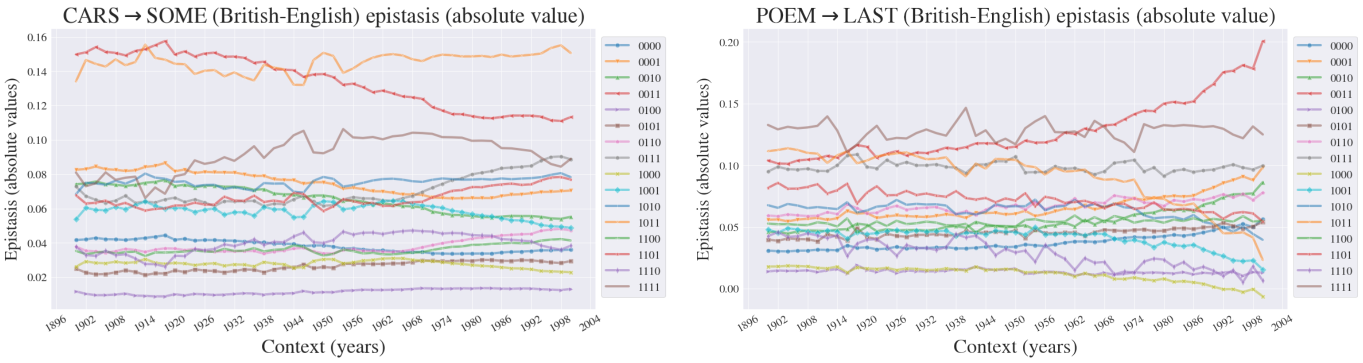


**(a)** **(b)**

**Figure S4: Dis-aggregated British-English higher-order epistasic effects.** As discussed in several places throughout this manuscript, dis-aggregated graphs represent how individual epistatic terms interact across contexts. These graphs represent those effects for the CARS → SOME and POEM → LAST landscapes in British-English.

#### Within-path competition ($C_w$) analysis British-English

Here we provide the results of an analysis of within-path competition among the British-English variant of the CARS → SOME and POEM → LAST landscapes examined in our primary study.

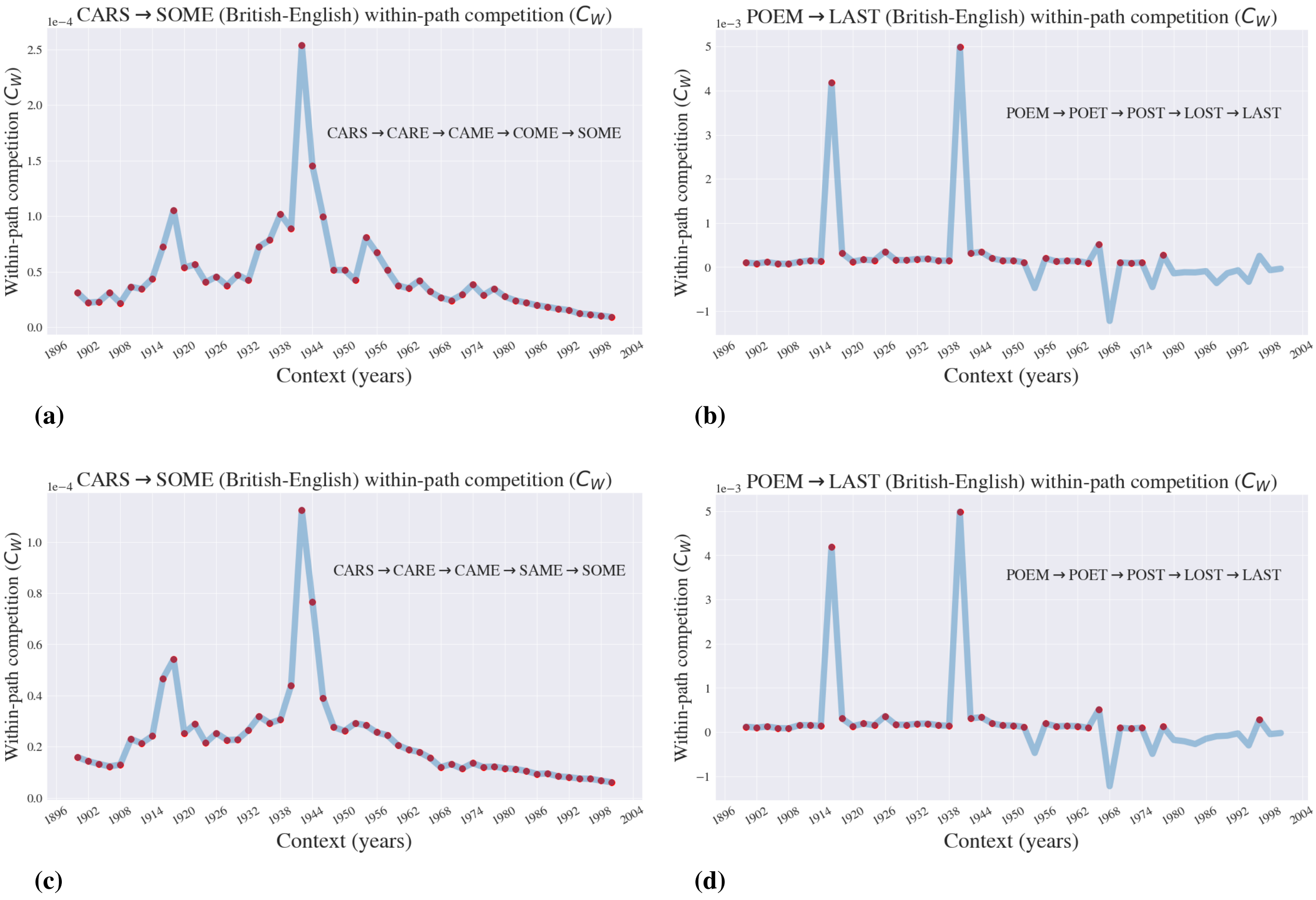


**Figure S5: British-English Within-path Competition.** Here we present within-path competition in the British English Lexical Landscapes subset

### Properties of 3 and 5 letter 1-grams

We have primarily focused our analysis on 4 letter 1-grams. This is not for any particular reason, and similar analyses could apply to landscapes of varying size. To demonstrate that the utility of Lexical Landscapes applies across word lengths, we provide the results of a complementary parallel investigation of random 3 and 5 letter 1-grams. We begin by presenting the fitness landscapes graphs for 3 and 5-letter examples: HAY → FIT and ADDED → VIRUS. The 3-letter landscape shows two uphill paths:
HAY → HAT → HIT → FIT and HAY → HAT → FAT → FIT. Then, we present the ($C_w$) coefficients for two paths, one in each landscape.

#### 3 and 5 letter fitness landscapes

In Figure S5, we observe the fitness graphs for HAY → FIT and ADDED → VIRUS.

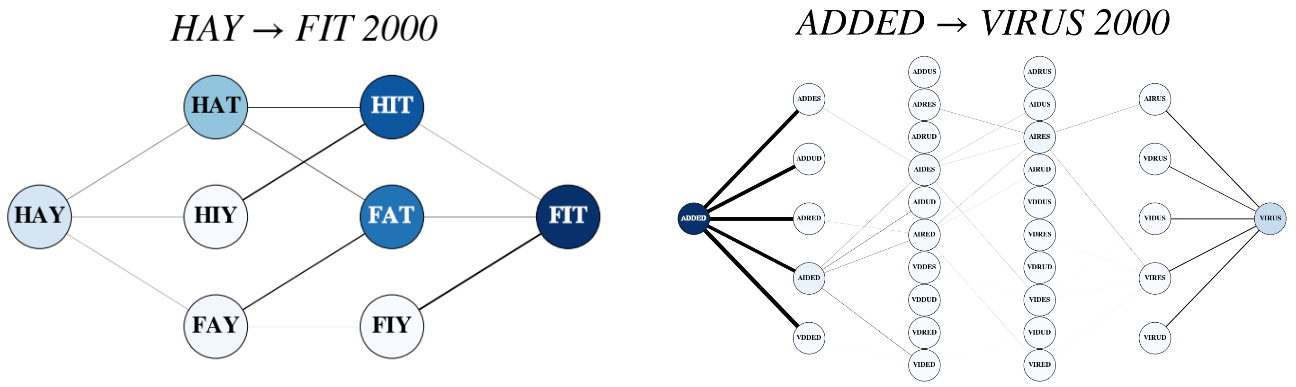


**(a)** **(b)**

**Figure S6: Fitness graphs for 3 letter (HAY → FIT) and 5 letter (ADDED → VIRUS) Lexical Landscapes.** Figures **(a)** and **(b)** are visualizations of the fitness landscapes for three and five letters. The color indicates the fitness: darker the blue, higher the fitness. Edges in the graph are weighted by the difference (in absolute value) between the fitness values of the two adjacent nodes and are emboldened in a proportional way to show the weight.

#### 3 and 5 letter within path competition ($C_w$)

Figure S6 describes the accessibility and

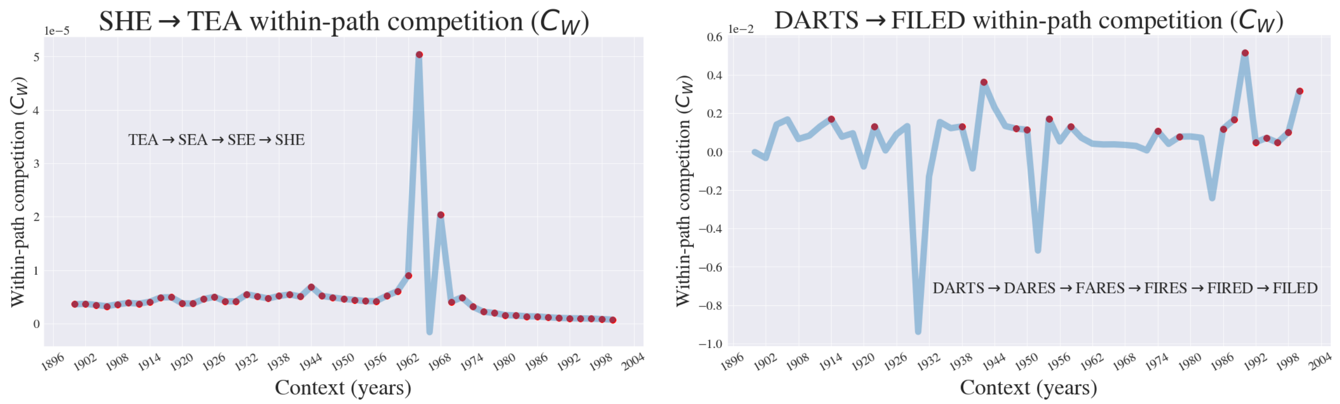


**(a)** **(b)**

**Figure S7: Within-path competition** ($C_w$) **for 3 and 5 world landscapes** Figures **(a)** and **(b)** demonstrate that the ($C_w$) can vary across environment (as it did for the 4 letter 1-grams discussed in the main texts). Red dots indicate the years where the specified path—shown to the right of each figure—represented an *uphill* path in the fitness landscape, that is, a path where each successive word in the path has a higher fitness than the previous one.

#### Epistasis dis-aggregated

Here we display the dis-aggregated epistasis coefficients for an example in the 3 and 5-letter word landscapes. Notice that in the 3-letter landscape there is little fluctuation in the epistatic coefficients, whereas the 5-letter landscape shows dramatic fluctuations over the years.

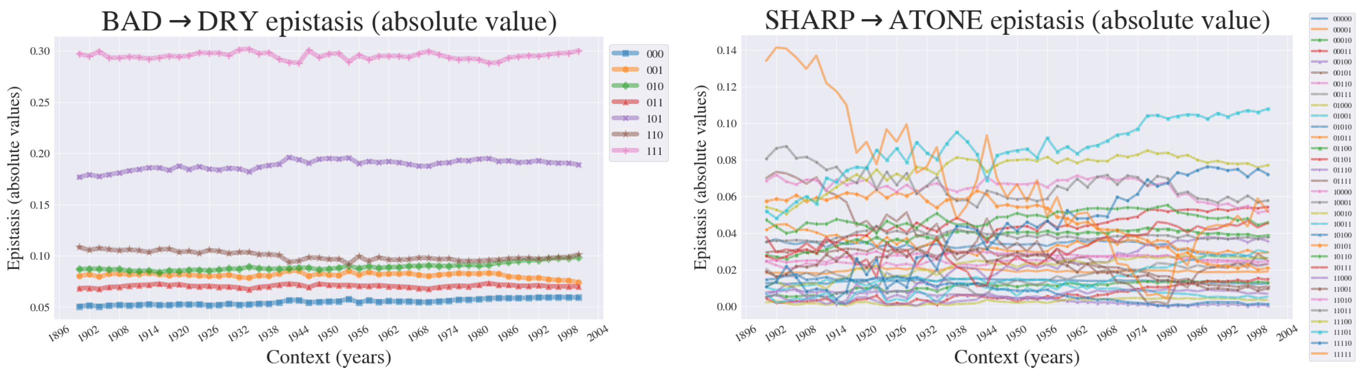


**(a)** **(b)**

**Figure S8: Epistasis across environment** Figures **(a)** and **(b)** show how epistatic effects for example three and five letter word landscapes can vary across environment, or in this case across time. We have chosen a three letter landscape (BAD to DRY with mild fluctuations over time to contrast it with the relatively large fluctuations in the five-letter landscape (SHARP to ATONE).
